# Actomyosin controls planarity and folding of epithelia in response to compression

**DOI:** 10.1101/422196

**Authors:** Tom P. J. Wyatt, Jonathan Fouchard, Ana Lisica, Nargess Khalilgharibi, Buzz Baum, Pierre Recho, Alexandre J. Kabla, Guillaume T. Charras

**Author notes:** co-first authors (Tom P. J. Wyatt, Jonathan Fouchard). Corresponding authors: Buzz Baum, Alexandre Kabla and Guillaume Charras.

## Abstract

Throughout embryonic development and adult life, epithelia are subjected to external forces. The resulting deformations can have a profound impact on tissue development and function. These include compressive deformations which, although hard to study in model systems due to the confounding effects of the substrate, are thought to play an important role in tissue morphogenesis by inducing tissue folding and by triggering mechanosensitive responses including cell extrusion and cell differentiation. Here, using suspended epithelia, we are able to uncover the immediate response of epithelial tissues to the application of large (5-80%) in-plane compressive strains. We show that fast compression induces tissue buckling followed by active tissue flattening which erases the buckle within tens of seconds. Strikingly, there is a well-defined limit to this second response, so that stable folds form in the tissue for compressive strains larger than ∼35%. Finally, a combination of experiment and modelling shows that the response to compression is orchestrated by the automatic adaptation of the actomyosin cytoskeleton as it re-establishes tension in compressed tissues. Thus, tissue pre-tension allows epithelia to both buffer against fast compression and regulate folding.

## INTRODUCTION

Epithelial tissues are frequently subject to in-plane compression, during both adult life and embryonic development, as the result of both intrinsic and extrinsic forces [1-4]. These forces are central to the function of many organs and are crucial for generating the complex shapes of adult tissues during developmental morphogenesis [5-7]. For example, in the airway, epithelia are subjected to periodic area changes as part of their normal physiological function during the breathing cycle and to longer term compression during diseased states such as asthmatic bronchial contraction [8; 9]. During embryonic development, compression guides a number of morphogenetic events involving tissue bending and folding, such as optic cup formation [7], villi formation in the gut [6], and the formation of cortical convolutions in the brain [10].

Recent work has revealed that epithelia have evolved a variety of cellular-scale mechanisms to detect and respond to compression via mechanotransduction [11]. For example, compressive deformations have been reported to activate signalling pathways that regulate cell differentiation, such as the nuclear relocalisation of β-catenin occurring during both Drosophila germband extension [12] and avian skin organogenesis [4]. Moreover, cultured epithelial monolayers can respond to the compression induced by cell crowding by triggering the extrusion of live cells in a process involving the mechanosensitive ion channel Piezo-1 [13]. Similar responses to crowding have also been observed *in vivo* in the Zebrafish tail fin and the Drosophila notum [13; 14].

Thus, compressive strain can lead to cell crowding or epithelial folding and, through mechanical signalling, to cell differentiation and cell extrusion. However, the mechanical properties which underlie these behaviours are not fully understood and the signalling pathways involved have not been clearly defined. This is in part because simple model systems have not been available to enable the evolution of the mechanical state of an epithelium to be studied following a compressive strain. This is especially the case for the immediate responses of epithelia to compressive strains.

To investigate the relationships between cell and tissue shape and stress during uniaxial in-plane compressive strain, we use suspended epithelia devoid of a substrate, allowing us to study the intrinsic response of the cells that make up the tissue in isolation from extracellular matrix (ECM). Using this approach, we find that epithelia can rapidly accommodate compression up to a well-defined limit of 35% compressive strain, which we term the buckling threshold. Before this threshold, actomyosin-generated tissue pre-tension allows the tissues to retain a planar morphology during slow compressive strains and to erase buckles induced by faster strains within tens of seconds. Tissue tension decreases linearly with compressive strain until approaching zero at the buckling threshold, at which point stable folds can be formed in the tissue. Importantly, the full range of observed tissue behaviours can be predicted by modelling epithelia as pre-tensed visco-elastic sheets which exhibit a buckling instability upon entering compression. Finally, we show that the buckling threshold is given directly by the ratio between actomyosin pre-tension and the tissue elasticity. These results provide a new framework for understanding how epithelial cells respond to compressive strains and tissue buckling.

## RESULTS

### Epithelia rapidly erase buckles in response to compressive deformation but stably buckle at high deformation

A major obstacle to understanding the response of epithelia to compression is the coupling between cells and their substrates, which confounds analysis in many *in vivo* and *in vitro* settings. The dominant mechanical properties of the substrate obscures stress evolution in the cellular part of the tissue and imposes the overall deformation. Therefore, to investigate the cell-intrinsic response of epithelial cells to compressive strains, we used cultured MDCK epithelial monolayers devoid of a substrate [15-17] (Fig. 1a).

**Figure 1.**
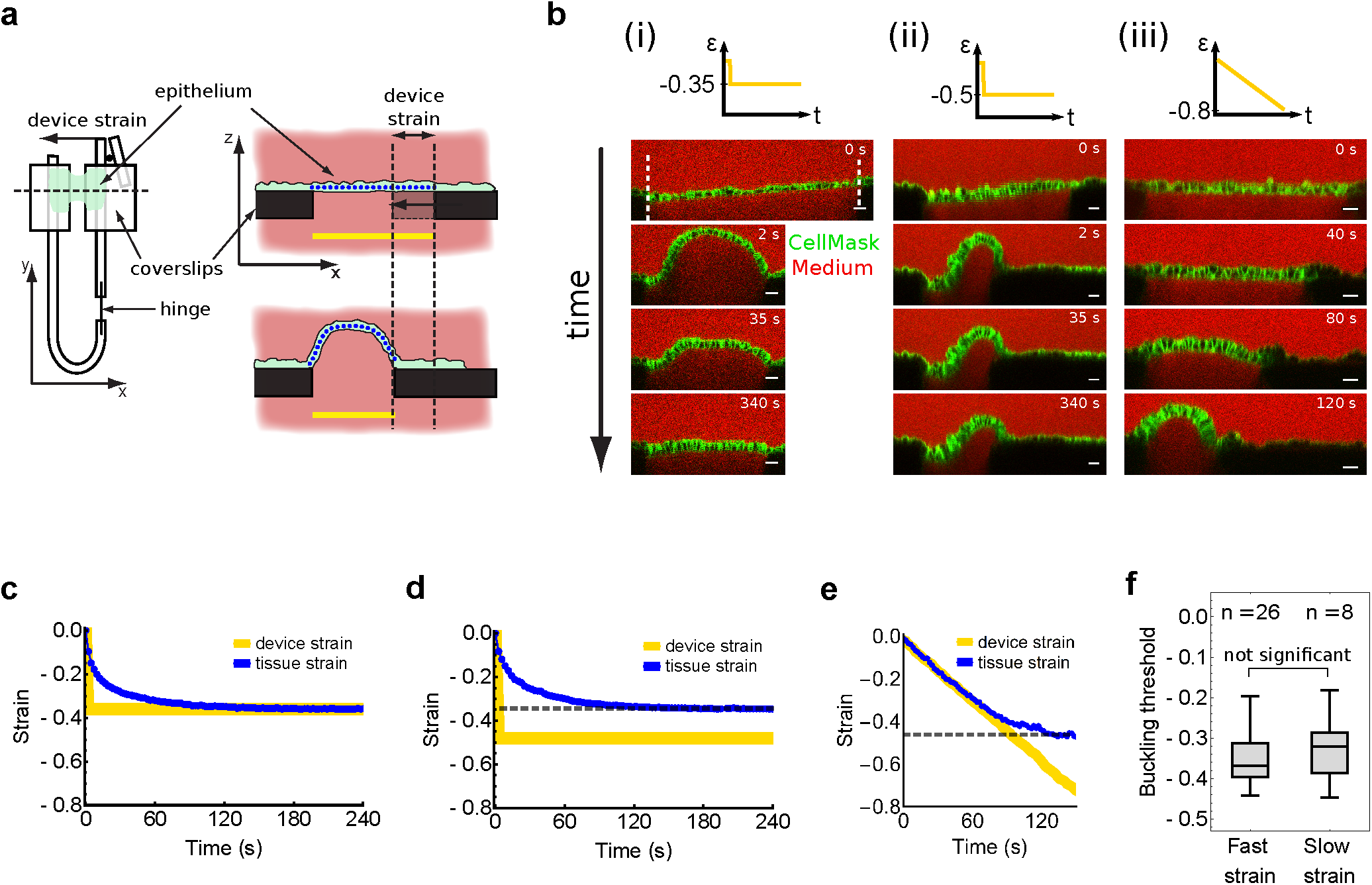
Epithelia rapidly erase buckles in response to compressive deformation but stably buckle at high deformation. **a** – Schematic diagram describing the method for application of compressive strain to suspended epithelia. Left: Top view of the device for mechanical manipulation of epithelial monolayers. An epithelial monolayer devoid of ECM (green) is attached between two coverslips affixed to the device. Uniaxial mechanical strain can be applied to the tissue by displacing the flexible arm with a motorized micromanipulator (black circle). Right: profile views of the setup along the dashed line in the left-hand diagram. Top: An epithelial tissue (green) is suspended between two coverslips (black). Bottom: the right hand-side coverslip is displaced and the tissue is deformed. The yellow line denotes the separation between the coverslips which allows calculation of the device strain. The blue dotted line denotes the midline of the tissue along which the tissue contour length is measured. **b** – Time series of profiles of MDCK epithelial monolayers before and during application of various compressive strains ε applied at different rates. Diagrams in the top row graphically represent the strain applied as a function of time. The white dashed lines indicate the points of attachment of the suspended monolayer to the coverslips. This delimits the region of the tissue upon which strain is applied. Strain was applied either as a step (*i-ii*) or as a ramp (*iii*). *i*: Step of intermediate strain (−35%) applied at high strain rate. *ii*: Step of large strain (−50%) applied at high strain rate. *iii*: Ramp of strain applied at low strain rate. Cell membranes are marked with CellMask (green), the medium is visualised by addition of dextran Alexa647 (red) making coverslips appear dark due to dye exclusion. Scale bars, 20 μm. The timing is indicated in the top right-hand corner of each image. **c** – Temporal evolution of the tissueengineering strain (blue) resulting from the variation of its contour length following a 35% reduction of the distance between coverslips, here referred to as device strain (yellow) applied at a strain rate of 500 %.s^-1^. **d** – Same as c, except the amplitude of the device strain is 50%. The dashed line marks the buckling threshold ε_b_. **e** – Temporal evolution of the tissue strain (blue) during a ramp of device strain (yellow) applied at low strain rate (0.5 %.s^-1^). The dashed line marks the buckling threshold ε_b_. **f** – Comparison of the buckling threshold ε_b_ measured following a step of fast compressive strain (as in **d**) and a slow ramp of compressive strain (as in **e**). The number of monolayers examined is indicated above the bars for each boxplot.

It is well understood from classical mechanics that slender elastic materials subjected to large compressive strains can undergo bending after a critical point called buckling. Here, after a step of −35% strain (‘device strain’, see Fig. 1a) applied at high strain rate (−500 %.s^-1^), most tissues took an arched shape (Fig. 1B (i) and Fig. S1a, left), reminiscent of buckling in solid materials. Less frequently (32% of experiments), epithelia took on a more complex wave-like shape reminiscent of the second mode of buckling (Fig. S1a, right).

Remarkably, the buckles were not stable configurations of the tissue. Instead the epithelia rapidly erased the buckle, becoming entirely planar within ∼1 minute (Fig. 1B (i) and Video 1). The evolution of global tissue strain was quantified by extracting the contour length of the tissue cross-section from live videos and comparing this to the contour length before strain application (see methods and Fig. S1b). Following the application of a step of compressive strain, the tissue strain first decreased at a very rapid rate before gradually slowing as the tissue approached a planar configuration (Fig. 1c). The average half-life of the flattening process was 4.4 ± 3.1 s (n = 17) (see methods).

HaCaT human keratinocytes, which we cultured as multi-layered epithelia, showed the same behaviour (Fig. S1c-e, Video 2), indicating that the response is not specific to a particular cell type or even tissue architecture and may be a generic property of epithelia.

To define the limit of this adaptive process, we applied a larger compressive strain at the same rate. The tissues exhibited the same initial behaviour, but could not completely accommodate the larger deformation (Fig. 1b (ii), Fig. 1d and Video 3). As a result, a buckle remained which was stable for periods greater than 10 minutes (Fig. S1f). The reduction of the tissue contour length after compression applied at high strain rate saturated at a strain of −34 ± 8 % (n = 26, Fig. 1f).

The existence of such a threshold suggests that a reference length of the tissue – independent of time – may have been reached. To test this hypothesis, the global tissue strain was measured during a large compression applied at a much lower strain rate (−80% at −0.5 %.s^-1^). Here, the monolayer maintained a planar morphology for large deformations until taking on an arched shape, at which point its length did not evolve any further (Fig. 1b (iii), Fig. 1e and Video 4). The maximum reduction of tissue contour length during these experiments was −33 ± 8 % (n = 8), which is not statistically distinguishable from the maximum deformation attained after fast application of compression (p = 0.45, Fig. 1f).

To further test that the same steady state deformation was reached by the tissue regardless of the history of deformation, we applied both a fast step and a slow ramp of strain to the same samples, separated by a 6 minute rest period. The ratio between the maximum deformations reached in the two cases was not distinguishable from 1 (p = 0.54, t-test, Fig. S1g). This enables us to define a buckling threshold ε_b_, above which the monolayer can no longer decrease its length.

These findings show that epithelial tissues rapidly adapt to large in-plane compressive strains by reducing their length, either restoring or maintaining a planar morphology, up until a well-defined limit, the buckling threshold, that is independent of the history of deformation.

### Tissue flattening is achieved through cell shape change and requires actomyosin contractility

To identify the mechanisms responsible for the tissue response to compression, we first examined the response at the cellular scale. During developmental morphogenesis, combinations of cellular behaviours, such as divisions, extrusion events, neighbour exchanges, and cell shape changes underpin changes in tissue shape and size [3].

Cell extrusions, cell divisions and cell-cell neighbour exchanges, which typically occur at a time-scale of tens of minutes, could not account for tissue flattening in our system which occurred at a time-scale of tens of seconds. In line with this, when we compared images of the cell-junction network before and after compression, it was clear that the deformation could be recapitulated by applying an affine deformation to the network prior to compression (Fig. S2a), indicating that changes in cell shape could account for the majority of the changes in tissue shape. Similarly, when we measured the shape of individual cells in planar tissues after the application of compressive strains of increasing magnitudes, the decrease in cell length along the axis of applied compressive strain (x) closely matched the tissue deformation, while the length remained unchanged along the perpendicular (y) axis (Fig. 2a and b) and cell height (z) increased. In sum, the total change in cell shape fitted a model of constant cellular volume (Fig. 2b, dashed lines and S2b). In accommodating the imposed compressive strain, cells therefore underwent a transition from a cuboidal to a more columnar shape, which accounted for the change in tissue shape.

**Figure 2.**
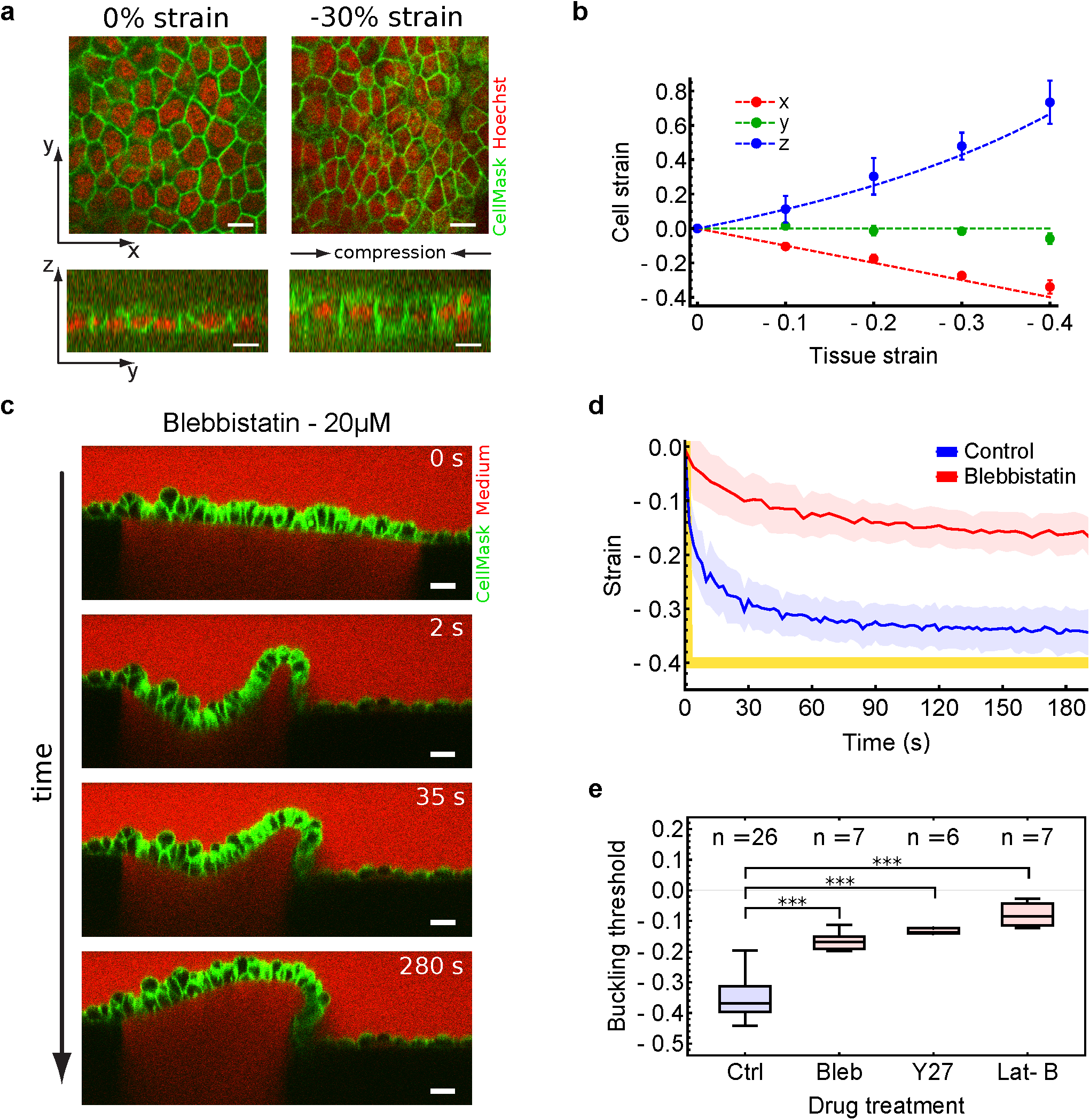
**Tissue flattening is achieved through cell shape change and requires myosin II contractility**. **a** – Confocal images of MDCK epithelial monolayers before (left) and during (right) 30% uniaxial compressive strain. Plasma membranes are marked with CellMask (green) and nuclei with Hoechst-33342 (red). Top: Single x-y confocal optical section through the middle of the monolayer. Bottom: y-z profile. Compressive strain is applied along the x-axis. Scale bars, 10 μm. **b** – Cellular strain as a function of the strain applied to the tissue. Solid circles denote cellular strain along the x-axis (red), y-axis (green), and z-axis (blue). Error bars denote standard deviation. Dashed lines indicate the predicted cellular strain assuming cell strain accounts entirely for tissue strain and that cells maintain constant volume. **c** – Profile of an MDCK monolayer treated with blebbistatin (20 μM) and subjected to fast compressive strain. Scale bars, 20 μm. Time is indicated in the top right-hand corner. **d** – Temporal evolution of tissue strain (mean ± SD) following fast compression for control (blue) and blebbistatin treated (red) tissues. Device strain is depicted by the yellow line. **e** – Buckling threshold ε_b_ inferred from the maximal tissue strain reached after fast compression. Tissues were treated with drugs perturbing actomyosin contractility. Bleb = Blebbistatin (20 μM); Y27 = Y-27632 (10 μM); Lat-B = Latrunculin B (3 μM). *** denotes statistically significant difference, p < 0.001. The number of monolayers examined in each condition is indicated above the bars for each boxplot.

Changes in tissue and cell shape could result from the relaxation of the viscous stresses which may have caused the transient surpassing of the critical buckling stress. Thus, since actomyosin activity is known to drive both cell shape changes of mammalian cells [18] and stress relaxation in single cells [19], we explored the role of actomyosin in this process. We therefore repeated the compression experiments after treating the tissues with a range of pharmacological inhibitors of actomyosin activity. While control tissues buckled upon fast compression as observed above, the flattening process was severely compromised by inhibitors of actomyosin contractility (Blebbistatin, Y-27632, Latrunculin-B, see methods) (Fig. 2c,d and Fig. S2c,d, Video 5). Firstly, consistent with the reported role of actomyosin in driving stress relaxation, the rate of flattening was reduced by inhibition of actomyosin contractility (Fig. S2e,f). Secondly, and more surprisingly, the tissue’s ability to accommodate strain saturated at smaller values upon inhibition of actomyosin (Fig. 2e). Importantly, this suggests that the buckling threshold itself depends on actomyosin. In the case of blebbistatin treatment (20 μM), for example, the buckling threshold was reduced to-17 ± 4 % (n = 7).

These results suggest that actomyosin activity not only permits the rapid adaptation of epithelia to compressive strain but also, through regulating the buckling threshold, regulates the transition between planarity and folding over prolonged compressive deformations.

### An actomyosin generated tissue pre-tension buffers against compression to prevent stable buckling of epithelia

While buckling is known to occur under compressive stress in inert materials, epithelial tissues are often naturally in a state of pre-tension originating from the actomyosin cytoskeleton [20-22]. We therefore hypothesised that actomyosin activity could tune the buckling threshold by controlling the amount of pre-tension in the epithelium. To test this hypothesis, we first aimed to determine the level of stress borne by the suspended epithelia before it was deformed. To this end, we prepared the tissues on a force sensitive device, consisting of two test rods: one flexible (serving as a force sensor) and the other rigid (Fig. 3a and methods). Tissue pre-stress was quantified by comparing the positions of the flexible wire before and after detachment of the monolayer from the device. The pre-stress in control conditions was tensional with a magnitude of 280 ± 40 Pa (n = 15) (Fig. 3b) - close to that typically measured in single cells [23] and dropped dramatically when actomyosin contractility was perturbed (Fig. 3b). This confirms that the generation of pre-tension requires actomyosin and that the reduction in buckling threshold caused by actomyosin perturbation (Fig. 2e) occurred in the context of lower tissue pre-tension.

**Figure 3.**
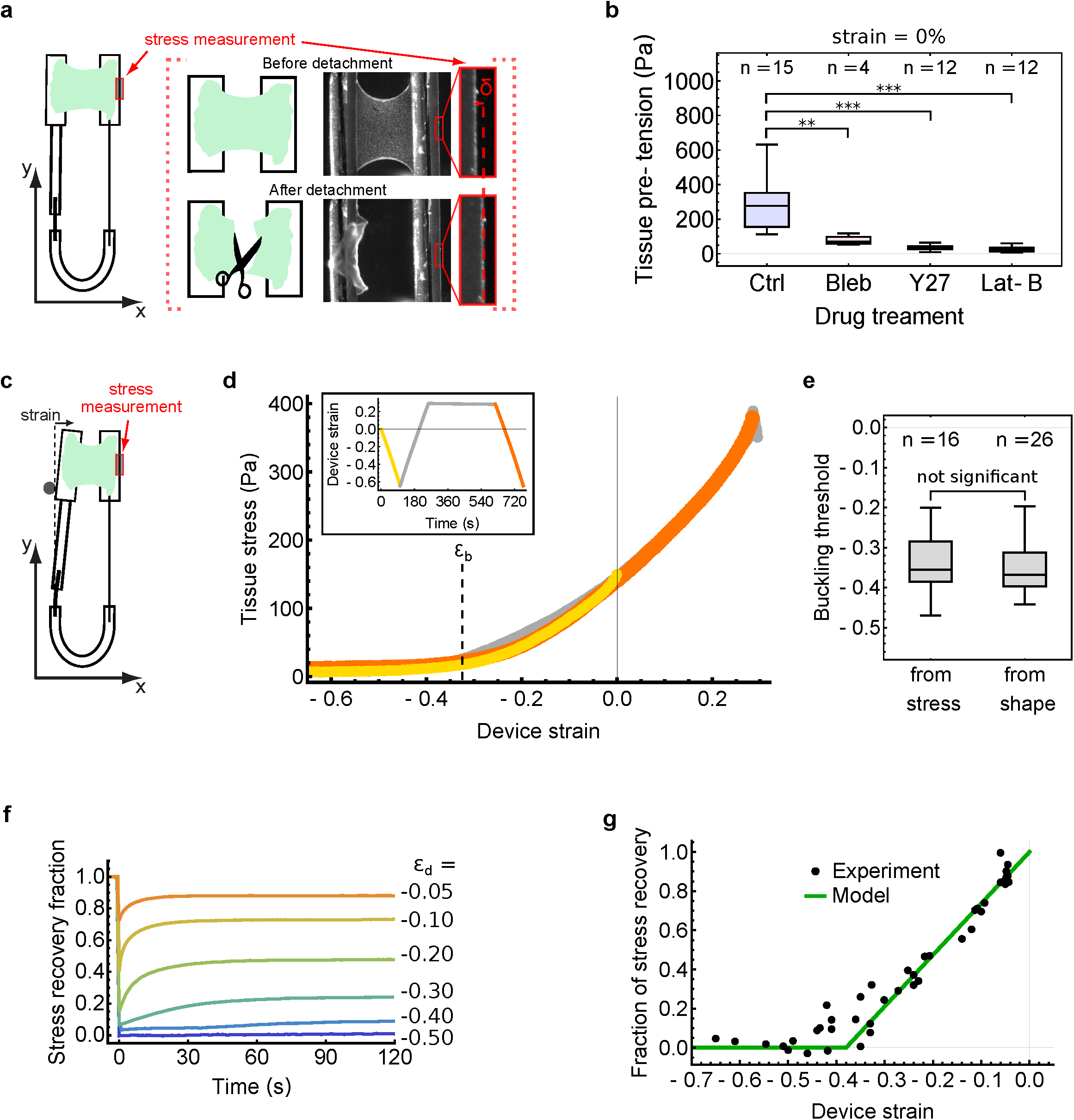
Tissue pre-tension buffers against compressive strain to prevent stable buckling of epithelia. **a** – Schematic diagram depicting the principle of measurement of tissue pre-stress. The monolayer is cultured between the two rods of the device (left). The left rod is the reference rod, while the right rod is flexible. Deflection of the right rod allows measurement of stress. The deflection of the flexible rod, δ (see enlarged region, furthest right), caused by tissue tension is measured from brightfield images acquired before (top) and after (bottom) detachment of the monolayer. **b** – Pre-stress of MDCK monolayers (i.e tissue stress at 0% strain) under various treatments inhibiting actomyosin contractility. ** denotes statistically significant difference, p < 0.01 and *** denotes p < 0.001. **c** – Schematic diagram for measurement of stress during application of compressive strain. The device is the same as in (**a**) but uniaxial strain is applied by displacing the left arm. Stress is measured at each time point by monitoring the deflection of the right arm. The rest position is determined by detaching the monolayer at the end of the experiment as shown in (**a**). **d** – Tissue stress as a function of the applied device strain during deformation at low strain rate (0.5 %.s^-1^). Inset: Time course of the strain applied with the device. The stress follows the same trend before (yellow) and after (orange) a period of 6 minutes of 30% stretch. The dashed line indicates the buckling threshold ε_b_. **e** – Comparison of the buckling threshold as measured from the transition identified in stress-strain curves (as in 3d) and from the maximum strain identified during imaging experiments (as in Fig. 1f). **f** – Time evolution of stress after application of compressive strains of various amplitudes applied at high strain rate (100 %.s^-1^). The stress is normalised to the pre-tension in the tissue before deformation. The magnitude of device strain ε_d_ is shown on the right. **g** – Fraction of stress recovery at steady state as a function of device strain for deformation applied at high strain rate. The green line shows the behaviour predicted by the rheological model presented in Fig. 4. Measurements from n = 42 monolayers. In boxplots (**b**, **e**), the number of monolayers examined is indicated above the bars.

To characterise how tissue stress evolved during compressive strain we applied a slow deformation to the monolayers (−0.5 %.s^-1^, as in Fig. 1e) whilst measuring the tissue-level tension (Fig. 3c and methods). Tissue stress initially decreased linearly with strain, before transitioning, at a stress of 28.8 ± 28.2 Pa (n = 16), to a second phase in which stress decreased with a smaller slope towards a plateau of zero stress (Fig. 3d, yellow). Such a stress-strain curve is a typical signature of a thin elastic sheet with a small bending modulus experiencing a buckling instability [24]. In support of this idea, the transition between the 2 phases occurred at a device strain of −33 ± 8 % (Fig. 3d, dashed line, and Fig. 3e, n = 16), which was indistinguishable from the buckling threshold identified previously from measurements of tissue shape during compression (p = 0.53, Fig. 3e). In addition, this transition point remained stable over multiple cycles of compression (Fig. S3a,b) or when a period of stretch preceded compression (Fig. 3d, orange). This further suggests that the buckling threshold is an intrinsic mechanical feature of the tissue.

Next, we investigated how the initial tissue stress evolved in response to rapid steps in compressive strain of various magnitudes (2-65%, as in Fig. 1c,d). For all ε, the stress decreased immediately upon application of strain, however, for all but the largest strains, stress was rapidly re-established in the tissue (Fig. 3f). The stress then plateaued at a level which depended on the magnitude of compressive strain applied, with a larger fraction of the pre-tension being recovered in tissues subjected to smaller strains (Fig. 3f). The curve relating steady-state stress to applied compressive strain for fast application of strain closely matched the stress response observed during slow applied strain (compare Fig. 3g and 3d). Indeed, the fraction of recovered stress decreased linearly with increasing compressive strain amplitude up to a transition point at a device strain of ∼-33%, below which there was no stress recovery.

Thus, the steady state stress in MDCK monolayers evolves linearly with compressive strain, independently of the history of deformation, and planar tissue morphologies can only be attained when this steady state stress is tensional. The generation of tensile stress is dependent on actomyosin contractility which buffers against compressive strains and buckling. Taken together, these data provide an explanation for the observed reduction of the buckling threshold upon perturbation of actomyosin activity (Fig. 2e).

### The response of epithelia to compressive strain is recapitulated in a model of a pre-tensed visco-elastic material which buckles under compressive stress

In sum, our findings demonstrate that there exists a quasi-static regime in which the tissue behaves as a pre-tensed elastic sheet with negligible bending stiffness. Additionally, the time-scales of tissue flattening (Fig. 1) and tension recovery (Fig. 3) suggest that a viscous contribution damps the response of epithelia during deformation. Consequently, we devised a simple rheological model consisting of a standard linear solid (SLS, in line with [15]) to which we added an active pre-tension (Fig. 4a and see Appendix 1 of supplementary materials). To account for the non-linearity at the buckling transition, we also supplemented the model with a ‘buckling condition’ in the form of a loss of tissue stiffness along the loading axis when the stress becomes negative (Fig. 4a).

**Figure 4.**
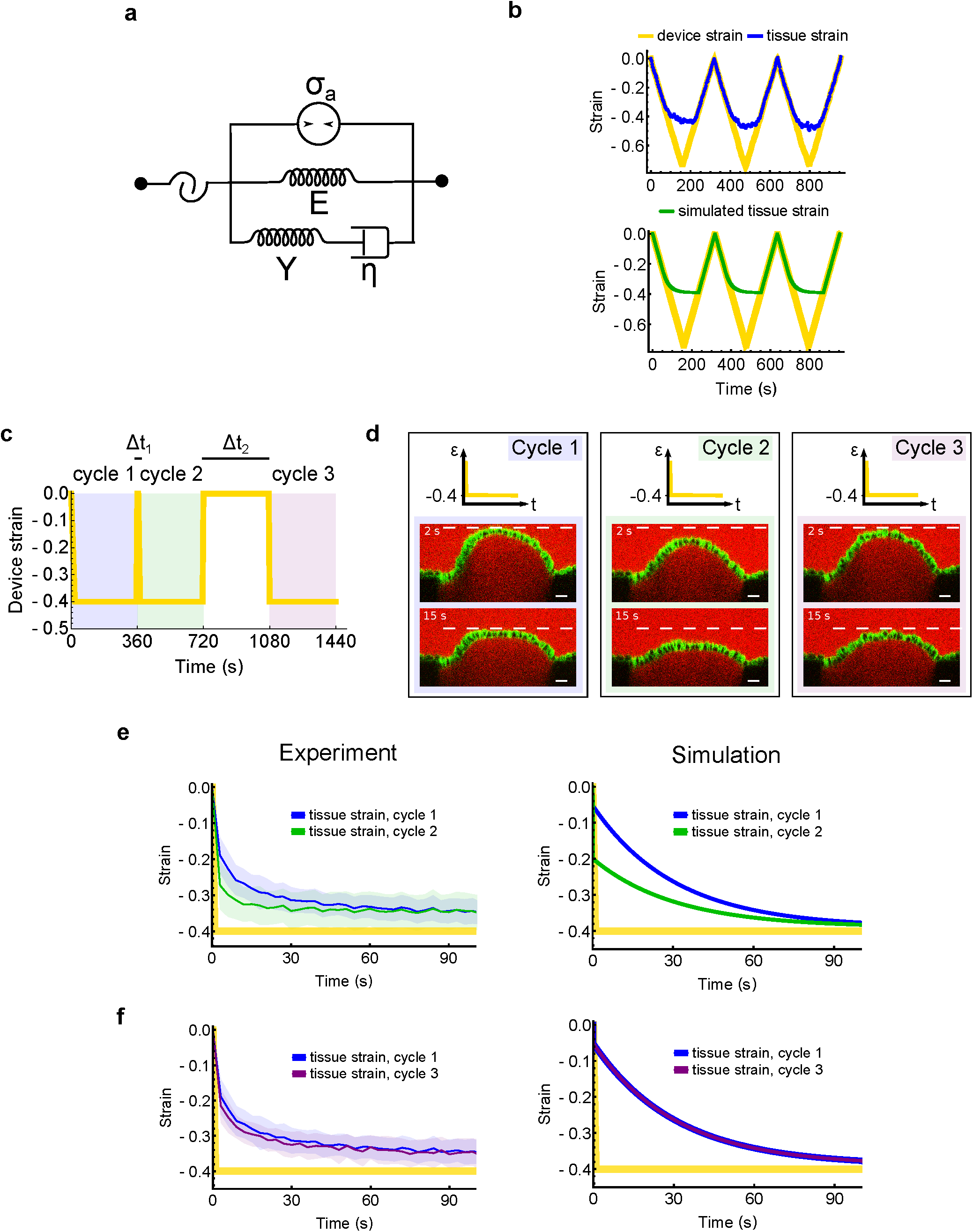
The response of epithelia to compressive strain is recapitulated in a model of a pre-tensed visco-elastic material which buckles under compressive stress. **a** – Diagram of the rheological model of the mechanical response of epithelia to compression. An active element (generating a stress σ_a_, top) models the contractile stress generated by myosin II. The middle spring (of stiffness E) models the elastic behaviour of the tissue at long time-scales, while the spring-dashpot element (of stiffness Y and viscosity η, bottom) models the short time-scale elastic behaviour and viscous relaxation. A condition of loss of stiffness is added to model the buckling instability (left): when the material reaches zero-stress, its stiffness is null. **b** – Temporal evolution of tissue strain in response to slow cycles of compressive strain (70% amplitude applied at 0.5 %.s^-1^). Top: representative experimental dataset. Bottom: model prediction. Device strain is depicted by the yellow line, experimental tissue strain is shown in blue, and the model prediction for tissue strain is shown in green. **c** – Sequence of device strain applied to the tissues to assess dependence of the response on strain history. Three cycles of 40% compressive strain were applied, each lasting 6 minutes. After each cycle the tissue was returned to its original length for a duration δt. Between cycles 1 and 2, Δt_1_ was 3 seconds and between cycles 2 and 3, Δt_2_ was 6 minutes. **d** – Profile images of MDCK monolayers shown 2 seconds and 15 seconds after compressive strain application. Diagrams in the top row graphically represent the strain applied as a function of time. Each column corresponds to a different cycle of compressive strain. For each time point, the dashed white line corresponds to the maximal vertical deflection of the tissue during the first cycle. Scale bars, 20 μm. **e** – Temporal evolution of the tissue strain (mean ± SD) after the first (blue) and second (green) cycle of compressive strain. In between these two cycles, the tissue was returned to its original length for Δt_1_ = 3 seconds, before being shortened again. Left: Tissue strain (mean ± SD) averaged over n = 12 experiments. Right: Model prediction. **f** – The same as (**e**) but for the first (blue) and third (purple) cycle of compressive strain. In between these cycles, the tissue was returned to its original length for Δt_2_ = 6 minutes, before being shortened again.

As observed in experiments, the assumptions of the model immediately imply that, upon application of compressive strain, the tissue may either buckle or remain planar, depending on the applied strain and strain rate (Fig. S4a and Appendix 1F of supplementary materials). For example, when deformation is applied at a sufficiently high strain rate, the tissue transiently buckles before reaching the buckling threshold (as in Fig. 1b (i)) because the spring-dashpot element behaves elastically and causes the tissue stress to decrease to zero.

To test whether the behaviour of epithelia subjected to compressive strain can indeed be captured by such a mechanical model, we extracted parameter values directly from experiments and simulated a set of mechanical perturbations. The active pre-stress, as shown in Fig. 3b, was σ_a_ = 280 ± 40 Pa. The stiffness of the elastic branch of the model E= 720 ± 50 Pa was extracted from the slope of the linear phase in slow compression experiments (Fig. 3d), while the stiffness of the viscous branch Y = 4770 ± 760 Pa and the relaxation time-scale τ = η / Y = 4.1 ± 0.7 s (with η the tissue viscosity) were extracted from a small subset of experiments in which stress recovery was measured in response to a small step of device strain (< 10%, Fig. S4b, see Appendix 2 of supplementary information for details). We refer to E as the long timescale stiffness, since this stiffness enables the formation of buckles which are stable for periods of > 10 min. Conversely, since Y only contributes to the stiffness at short time-scales, we refer to it as the short time-scale stiffness.

Firstly, the parameters σ_a_ and E directly predict a buckling threshold ε_b_ via the equation ε_b_ = -σ_a_ / E (see Appendix 1), giving a value of −39%, in agreement with experiments (Fig. 1f and 3e). This is also recapitulated in simulations of compressive strain applied at low strain rate for which the model accurately reproduces the experimentally observed global tissue strain (Fig. 4b). Secondly, the different regimes of stress recovery after fast compression associated to different amplitudes of compressive strain also emerged in simulated tests (Fig. 3e, 3f, Fig. S4c and Appendix 1 of supplementary materials).

### Viscous properties endow epithelia with reversible memory of past compression

A further prediction of the model was that, unlike the buckling threshold ε_b_, the time required for the tissue to flatten was dependent on the history of deformation (Appendix 1 of Supplementary Materials). To test whether this was the case experimentally, three cycles of 40% compressive strain, were applied to the tissue (Fig. 4c). After each application of compressive strain, held at each time for 6 minutes, the tissue was returned to its original length for a chosen duration Δt. Between the first and second cycle, Δt_1_ was just 3 seconds whereas between the second and third, Δt_2_ was 6 minutes.

The tissues reached a planar shape significantly faster after the second application of compressive strain compared with the first (Fig. 4d,e and Video 6), confirming that indeed the time to flatten does depend on the tissue’s history of deformation. This ‘memory effect’ was recapitulated in the model (Fig. 4e) and is due to the incomplete relaxation of the viscous element during the transient lengthening of the tissue.

Conversely, after 6 minutes rest at the initial length, which is sufficient for full relaxation (Δt_2_, Fig. 4a, 3^rd^ cycle), the time required to become planar was indistinguishable from that in the first compressive period. This demonstrates that the ‘memory’ of the past compression could be fully reversed with time (Fig. 4d,f and Video 7).

### Junctional F-actin determines the long time-scale stiffness of epithelia and cooperates with actomyosin contractility to regulate the buckling threshold

In sum, a broad range of steady-state and dynamic behaviours of the epithelial response to compressive strain can be fully recapitulated by modelling it as a pre-tensed visco-elastic material determined by just 4 parameters and endowed with a buckling condition. In our model, the buckling threshold of the monolayer ε_b_ emerges as the ratio between the pre-tension σ_a_ and the long time-scale stiffness E of the tissue. ε_b_ and σ_a_ are bothcontrolled by the actomyosin cytoskeleton (Fig. 2e and 3b). Therefore, to fully determine the molecular origins of the monolayer behaviour, we aimed to identify what controls the tissue stiffness E.

One key difference in the actomyosin organisation of isolated cells and epithelia is the presence of intercellular junctions. In epithelial monolayers, it has been proposed that at least two clearly distinct actomyosin networks coexist: junctional and cortical F-actin. The junctional network has slow turnover and appears associated with intercellular adhesions, while the cortical network has a faster turnover similar to that of cortical F-actin in isolated cells [25].

To test the importance of the junctional actin network in determining the long time-scale stiffness E, we subjected monolayers to prolonged Latrunculin B treatment at various concentrations and measured their stress-strain response at low strain rate. At the highest concentration (1μM), actin and cadherin staining revealed that the majority of the junctional F-actin network was disrupted (Fig. 5a) with only small patchy remnants of junctional F-actin structures (in agreement with [26]). A less pronounced loss of junctional F-actin was observed after treatment with 0.1 μM Latrunculin B (Fig. 5a).

**Figure 5.**
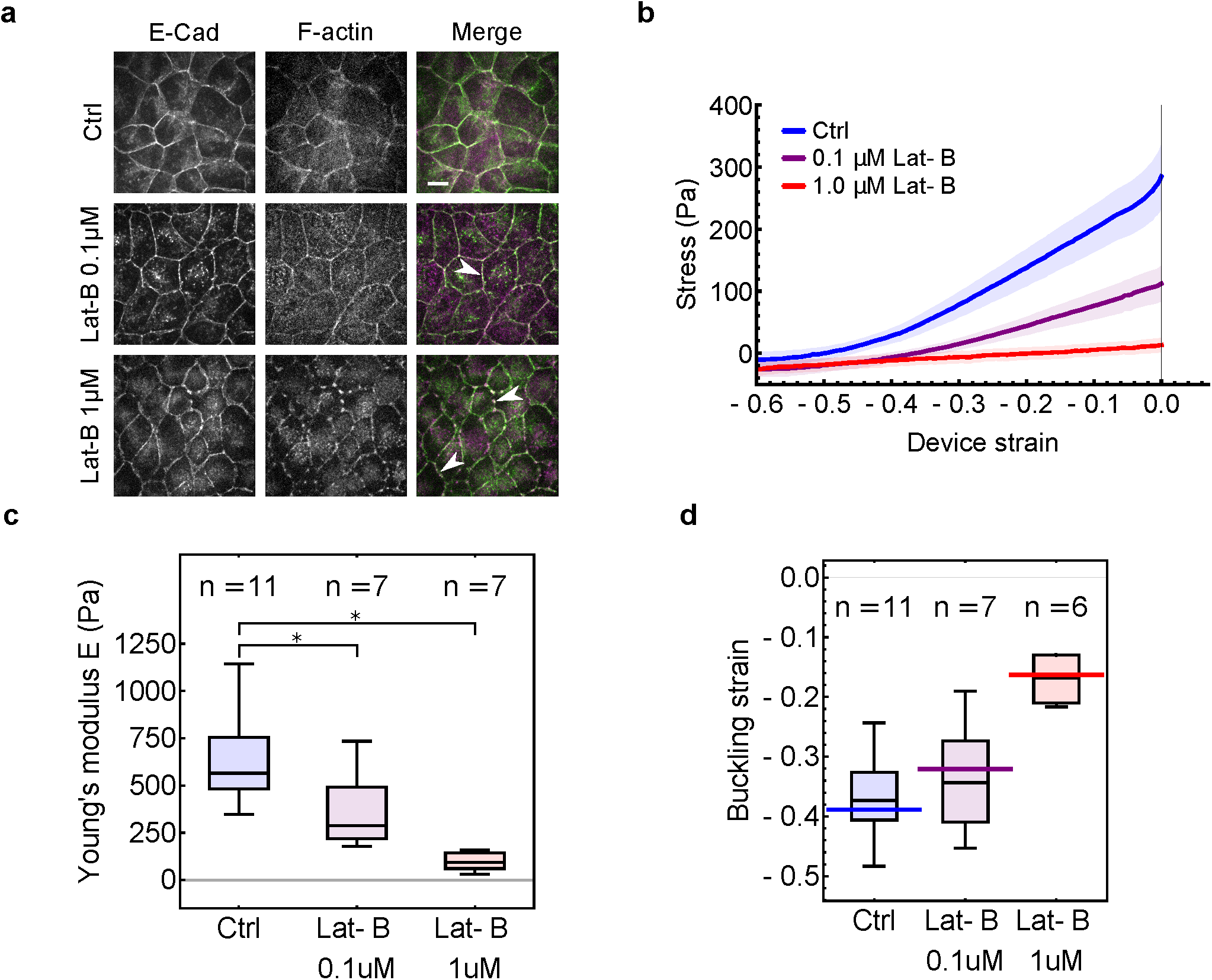
Junctional F-actin determines the long time-scale stiffness of epithelia and cooperates with actomyosin contractility to regulate the buckling threshold ε_b_. **a** – Effect of Latrunculin B treatment on intercellular junction organisation. MDCK ECadherin-GFP (green) cells were stained with phalloidin-Alexa647 to visualise F-actin (magenta). White arrowheads show regions where the junctional F-actin network is disrupted due to Lat-B treatment. Scale bar, 10 μm. **b** – Tissue stress measured during compressive strain applied at low strain rate (0.1 %.s^-1^). Control tissues are shown in blue, tissues treated with 0.1 μM Latrunculin B are shown in purple, and those treated with 1 μM Latrunculin B are shown in red. n ≥ 7 tissues for each treatment. Note that the strain rate has been chosen to ensure that Latrunculin B treated samples are tested in a quasi-static regime. **c** – Young’s modulus E (as a measure of long time-scale stiffness) of the tissue for control and Latrunculin B treated samples (0.1 μM and 1 μM). Young’s modulus was measured from the slope of the curve in (**b**) before transition to buckling. * denotes statistically significant difference, p < 0.05. **d** – Buckling threshold for control and Latrunculin B treated samples (0.1 μM and 1 μM) as measured from the transition in the stress-strain curve. The solid coloured bars correspond to the buckling threshold determined from the measures of Young’s moduli and pre-tension according to the rheological model shown in Fig. 4a. In boxplots (c, d), the number of monolayers examined in each condition is indicated above each box.

Stress-strain curves acquired during compressive strain application were similar to non-treated samples, consisting of a first regime where stress was proportional to strain, followed by a saturation around zero stress at high compressive strain (Fig. 5b). However, the slope of the first regime, the stiffness E, was affected by latrunculin treatment in a dose-dependent manner. At the highest concentration, E decreased ∼10-fold, suggesting that junctional F-actin is a major contributor to MDCK monolayer stiffness (Fig. 5c). Smaller concentrations of drug had an intermediate effect on stiffness, indicating that it can be adjusted smoothly by F-actin regulation (Fig. 5b,c).

Consistent with single cell measurements [23], when F-actin content was decreased, both pre-tension and tissue stiffness were decreased (Fig. S5). As predicted by our model, the buckling threshold was only mildly changed when both tissue stiffness and pre-tension were both lowered to a similar extent as a result of low concentration Latrunculin B treatment (Fig. 5d). However, for the highest dose of drug, while tissue pre-tension was almost entirely abolished, the significant residual tissue stiffness resulted in a large reduction in buckling threshold (Fig. 5d). Indeed, the buckling threshold (Fig. 5d, boxplots) closely matched that predicted by the model ε_b_ = -σ_a_ / E (Fig. 5d, solid lines).

These findings therefore demonstrate that the buckling threshold ε_b_ of epithelia is determined by the ratio between a myosin II-dependent pre-tension σ_a_ and the stiffness E controlled by junctional F-actin, which likely defines a network elasticity in the monolayer.

## OUTLOOK

Our results reveal that epithelia can accommodate remarkably large and rapid reductions in their surface area. This response to compressive strains arises from isometric cell shape changes and is orchestrated autonomously by the actomyosin cytoskeleton, which not only controls the dynamics of the mechanical response but also the transition from a planar to a folded morphology. The full range of behaviours observed could be reproduced by a simple zero-dimensional mechanical model of the epithelium in the form of a pre-tensed visco-elastic sheet which exhibits a buckling instability upon switching between a tensile and compressive state. The visco-elastic properties of the tissue set the time-scale of tissue flattening and cell shape adaptation. This time-scale – on the order of tens of seconds – is consistent with the time-scale of tissue relaxation observed in laser severing experiments in the *Drosophila* pupa dorsal thorax epithelium [27] and that of the equilibration of cell-cell junctions subjected to constant stretching force in the early *Drosophila* embryo [28; 29].

Our results also demonstrate that the tissue response to compressive strain possesses a well-defined limit at a strain of ∼-35%, i.e. the buckling threshold, above which stable folds can be formed in the tissue. The buckling threshold is regulated by the interplay between myosin II generated pre-tension and tissue elasticity which is controlled by junctional F-actin – the ratio of these two quantities defines the strain at which the tissue reaches compressive stress. The resulting buckling instability may act in parallel to other well-studied mechanisms of epithelial bending and folding, which include differential growth of connected tissues [6;30] and spatially patterned force generation driven by actomyosin contractions [31]. Indeed, apical (or basal) constrictions are often preceded *in vivo* by an increase in cell density before fold formation [2;7]. This implies that polarized constriction is accompanied by a planar compressive deformation of cells, which could help to robustly control tissue folding. Thus, future studies should consider a role for the reduction of tissue pre-tension during epithelial folding.

Together, the buckling threshold and flattening time-scale define the maximum strain and strain rate which can be imposed upon an epithelium before it becomes subjected to compressive stresses. Such stresses can be damaging to the cells [32] and stress accumulation at the epithelium-substrate interface can lead to delamination of the epithelium. The buckling threshold and flattening time-scale are therefore crucial material parameters for understanding the response of any epithelium subjected to compressive strain during morphogenesis or during the deformations that accompany normal organ physiology. On longer time-scales, this further underlines the need for cellular-scale mechanisms such as cell delamination in epithelia to counter the deleterious effects of prolonged compression [13; 14]. Finally, our results also raise the possibility that actomyosin pre-tension may play a role in various cell [33] and tissue contexts throughout evolution to act as a buffer against unwanted stresses and distortions of shape that may otherwise be caused by compression.

## ACKNOWLEDGEMENTS

The authors wish to acknowledge present and past members of the Charras, Baum, and Kabla labs for stimulating discussions. T.W. and N.K. were part of the EPSRC funded doctoral training program CoMPLEX. J.F. and P.R. were funded by BBSRC grant (BB/M003280 and BB/M002578) to G.C. and A.K. N.K. was funded by the Rosetrees Trust, and the UCL Graduate School. A.L. was supported by an EMBO long term post-doctoral fellowship. B.B. was supported by UCL, a BBSRC project grant (BB/K009001/1) and a CRUK programme grant (17343). T.W., J.F., N.K., A.L. and G.C. were supported by a consolidator grant from the European Research Council (MolCellTissMech, agreement 647186).

## AUTHOR CONTRIBUTIONS

T.W., J.F., B.B. and G.C. designed the experiments. T.W., J.F., A.L., and N.K. carried out the experiments. T.W. and J.F. performed the data and image analysis. P.R. and A.K. designed the rheological model. T.W., J.F., B.B. and G.C. wrote the manuscript. All authors discussed the results and manuscript.

## METHODS

### Cell Culture

MDCK and HaCaT cells were cultured at 37°C in an atmosphere of 5% CO_2_. Cells were passaged at a 1:5 ratio every 3-4 days using standard cell culture protocols and disposed of after 30 passages. For MDCK cells, the culture medium was composed of high glucose DMEM (Thermo Fisher Scientific) supplemented with 10% FBS (Sigma) and 1% penicillin-streptomycin (Thermo Fisher Scientific). MDCK-E-Cadherin-GFP cell lines (generated as described in [16]) were cultured in the same conditions as wild-type cells except that 250 ng/ml puromycin was added to the culture medium.

HaCaT cells were cultured in low calcium conditions, consisting of a minimal DMEM supplemented with 0.03 mM CaCl_2_, 10% calcium-free FBS, 1% penicillin-streptomycin and 1% L-Glutamine (Gibco).

### Fabrication of tissue mechanical manipulation and measurement devices

For profile imaging of epithelia during compression, tissues were cultured on the devices described in [17]. Briefly, stretching device arms were built from glass capillaries (Sutter Instruments) and a length of nickel-titanium (nitinol) wire (Euroflex) that acted as a hinge. Glass coverslips (VWR) that acted as a substrate for cell culture were glued to the glass capillaries. To allow precise control over both compressive and tensile strains, another small piece of glass capillary was added to the hinged side of the device at an angle to allow continuous contact with the micro-manipulator probe (see illustration in Fig. 1a).

The stress measurement devices were an adaptation of the force measurement device described in [16]. Briefly, a nickel-titanium (nitinol) wire was glued into a bent glass capillary. Then, tygon cylinders were glued to the extremities of the capillary and wire.Here, a hinge was added at the base of the rigid rod to control the applied deformation from this side while the force was measured by imaging movement of the flexible wire (see illustration in Fig. 3a).

### Generation of suspended tissues and preconditioning

Suspended monolayers of MDCK cells were generated as described in [16]. Briefly, a drop of collagen was placed between the test rods and left to dry at 37°C to form a solid scaffold. This collagen was then rehydrated before cells were seeded onto it and cultured for 48-72 hours. Immediately before each experiment the collagen scaffold was removed via enzymatic digestion. HaCaT monolayers were made using the same procedure. During the generation of HaCaT monolayers, standard culture medium (see above for MDCK) was used to allow the formation of robust intercellular junctions rather than the low calcium medium used for passaging the cells.

Before each experiment, the tissues were preconditioned by applying 5 cycles of 30% stretch at a rate of 1 %.s^-1^. The tissues were then left unperturbed for 6 minutes before application of compressive strain.

### Confocal imaging of tissues and mechanical manipulation

Tissues were imaged at 37°C in a humidified atmosphere with 5% CO_2_. The imaging medium consisted of DMEM without phenol red supplemented with 10% FBS.

To visualise the shape of the tissue during mechanical manipulation, cell membranes were labelled with CellMask membrane stain stained before imaging via a 10 minute incubation with 1X CellMask plasma membrane stain (Thermo Fisher Scientific). AlexaFluor-647-conjugated dextran, 10,000 MW (Thermo Fisher Scientific) was added at 20 μg.mL^-1^ to the imaging medium to visualise by exclusion the coverslips on which the epithelia were grown.

Profile views of the tissues during mechanical manipulation were obtained using a 30X objective (UPLSAPO S, NA=1.05, Olympus) mounted on an Olympus IX83 inverted microscope equipped with a scanning laser confocal head (Olympus FV1200). Each image consisted of roughly 200 slices spaced by 0.5 μm. Time series were with an interval of ∼2.2 seconds.

For imaging of cell shape change, cell membranes were visualized with CellMask. Z-stacks were acquired using a 60X objective (UPLSAPO, NA=1.3, Olympus) mounted on a spinning disk confocal microscope which consisted of a Yokogawa spinning disk head (CSU22; Yokogawa) and an iXon camera (Andor) interfaced to an IX81 Olympus inverted microscope.

To apply the mechanical deformation during confocal imaging, a custom-made adaptor was wedged in the top end of the hinged arm of the stretching device. The adaptor was connected to a 2-D manual micromanipulator mounted on a motorized platform (M-126.DG1 controlled through a C-863 controller, Physik Instrumente). The manual micromanipulators were used for initial positioning of the adaptor. Then, the tissues were deformed by moving the motorized platform via a custom-made Labview program (National Instruments).

### Quantification of device strain, tissue strain and flattening half-life

To precisely quantify the imposed compressive strain, the positions of the two coverslip edges which delimited the span of suspended tissue were determined by segmentation of dye exclusion in Alexa-647 fluorescence images. The strain imposed by the device was then defined as:

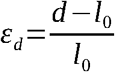

where *l*_0_ and *d* are the coverslip-to-coverslip distances before and after compressive strain, respectively.

To quantify tissue profile length evolution in response to compression, an implementation of the Chan-Vese algorithm [34] in Mathematica (Wolfram Research) was used to segment the tissues from the background. To convert each binary mask generated by segmentation into a line, morphological thinning was applied to the mask repeatedly until complete skeletonisation. The length of the shortest path between opposite edges of the image via foreground pixels thus constituted a measure of the cross-sectional contour length of the monolayer (see Fig. S1b). The tissue strain at each time point was defined as:

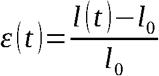

where *l*_0_ and *l* (*t*) are the suspended tissue length before deformation and at time t after compressive strain, respectively.

The half-life of tissue strain in response to a step of compression (Fig. 1) was defined as the time for the tissue to decrease its contour length by 50% of the total length change that occurred. This time was extracted after polynomial interpolation of the *ε* (*t*) data-points.

### Quantification of 3D cell shapes

The cell outlines were automatically segmented and hand-corrected using the Fiji [35] plugin *Tissue Analyzer [36]*. Measurements of cell shape were extracted from the resulting outlines using custom written routines in Mathematica (Wolfram Research). Cell length measurements were obtained by calculating the minimum bounding box of the cell outline. Bounding boxes were oriented along the experimental x-and y-axes with the x-axis corresponding to the axis of deformation. Measurements of cell height were obtained manually in Fiji from cross-sectional slices through the image stack along y-z planes.

### Drug treatments

Drug treatments were performed as follows. To block myosin contractility, Blebbistatin was added at a 20 μM concentration 10 minutes prior to experiments. To block Rho-kinase activity, Y-27632 was added at a concentration of 20 μM, 10 minutes prior to experiments. For complete depolymerisation of the F-actin cytoskeleton, we treated monolayers with 3 μM Latrunculin B for 30 minutes prior to experiments. For dose-dependent depolymerisation of junctional F-actin, we treated monolayers with either 1 μM and 0.1 μM Latrunculin B for 1 hour prior to experiments.

### Mechanical testing experiments

Mechanical testing experiments were performed at 37°C in Leibovitz’s L15 without phenol red (Thermo Fisher Scientific) supplemented with 10% FBS.

Tissues were imaged every second using a 2X objective (2X PLN, Olympus) mounted on an inverted microscope (Olympus IX-71). Images were acquired with a CCD camera (GS3-U3-60QS6M-C, Pointgrey). Compressive strain was applied using the motorized platform described above.

### Stress measurements during application of compressive strain

To measure the stress in the monolayer, we measured the displacement of the flexible arm of the stress measurement device from images before and after detachment of the monolayer (see Fig. 3a). The monolayers were detached from the reference arm of the device by cutting them with a tungsten needle. The force applied by the monolayer can be estimated by considering that the flexible arm acts as a cantilevered beam.

We define 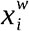as the position of the wire when the monolayer is attached between the barsand *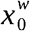*as the position of the wire when the monolayer is detached from the bars (the rest position). The force applied by the monolayer on the wire is given by:

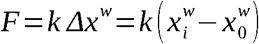

where k is the stiffness of the wire defined as:

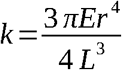

Here, *E* is the Young’s modulus of the wire, *r* its radius, and *L* its length. *E* was independently measured by loading wire with pieces of plasticine of different weights and measuring the deflection of the wire as described by (Harris, Nat. Protocols, 2014).

The stress of the tissue was then computed as:

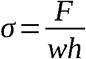

where *w* is the average width of the tissue determined from bright field images and *h* the tissue thickness. As shown in Figure 2a-b, tissue thickness was dependent on the applied strain. Previous work [15] and Figure 2b show that the cell volume remains constant during changes in monolayer area. Furthermore, past the buckling threshold (ε_b_∼0.4), the tissue length does not change and therefore the thickness also does not change. With these assumptions, we could estimate the evolution of the thickness *h* for calculation of *σ* as follows:

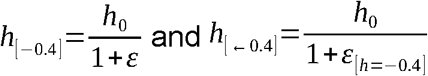

In experiments, the deflection of the wire *δ Δ xw* was determined by image cross-correlation with a sub-pixel accuracy through a custom-written algorithm based on the *register_translation* function of scikit-image, a Python image processing toolbox [37]. For the analysis, a portion of the wire was cropped from the images acquired during the experiments. The average error of the stress measurement method was measured from simulated displacements over 10 different samples. This gave 22 ± 31 nN (±SD) and 1.1 ± 1.6 Pa according to the average shape of the tested samples.

Tissue pre-stress measurements (*σ*_*a*_) were then obtained from the stress value before any deformation was imposed to the tissue, i.e *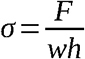*

For experiments where monolayer strain was varied, the position of the rigid rod was used to compute the displacement applied to the tissue. This position was extracted using the same method as for estimating the position of the flexible wire in stress measurements (see above). The device strain was defined as:

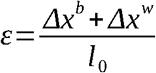

where 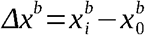refers to the displacement of the rigid rod, *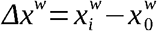* and *l*_0_ is the initial rod-to-rod distance.

Note that the deflection of the flexible wire *Δx^w^* due to the evolution of tissue stress over time was small compared to the deflection imposed by the movement of the rigid bar *Δx^b^*. For example, for a dataset where a deformation of −54 ± 5% was applied, the movement of the wire led to a variation in deformation of 6 ± 4% over the course of the experiment.

### Determination of the buckling threshold from stress-strain curves

Stress-strain curves were biphasic with stress first decreasing linearly with strain before saturating at high compressive strain. The transition between the two stress regimes was determined as follows. For each strain ε_i_ between [0, ε_max_], we fitted a linear function to the stress σ over the interval [0, ε_i_] and a constant over the interval [ε_i_,ε_max_]. For each ε_i_, the average of the sum of residuals is computed:

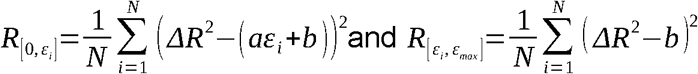

The transition point is then defined as the strain ε_i_ which minimizes the sum of the average errors over each interval:

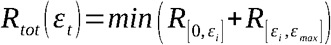

The stress at transition was then equal to σ_t_= σ(ε_t_). In addition, the elasticity of the tissue E could be extracted from the slope of the linear portion of curve.

### F-actin staining

MDCK Ecadherin-GFP cells were cultured to 90% confluence on glass substrate. They were fixed with 4% paraformaldehyde diluted in PBS for 20 minutes at room temperature, permeabilised in 0.2% Triton-X in PBS for 5 min and blocked with a solution of 10% horse serum in PBS for 30 min. To label F-actin, the cells were incubated with a solution of Phalloidin-Alexa 647 diluted at 1:40 from a stock solution of 200 units/mL for 1 hour at room temperature. The samples were then imaged on an Olympus FV-1200 confocal microscope.

### Date availability

The data that support the findings of this study are available from the corresponding author upon reasonable request.

## Other statistical and data analysis

All other routine data and statistical analysis were performed using the Python language environment and its scientific libraries (NumPy, SciPy) as well as Mathematica. Image processing was carried out with the Fiji package. All code is available from the corresponding author upon reasonable request. All boxplots show the median value (central bar), the first and third quartile (bounding box) and the range (whiskers) of the distribution. All tests of statistical significance are Mann-Whitney U tests unless otherwise stated. Measured values are given as mean ± standard deviation unless otherwise stated. Each dataset is pooled across experiments which were performed on at least 3 separate days.

## VIDEO LEGENDS

### Video 1

Buckling and flattening of an MDCK epithelial monolayer in response to a −35% compressive strain applied at high strain rate (500 %.s^-1^). Cell membranes are marked with CellMask (green), the medium is marked with Dextran Alexa647 (red) to allow for visualisation of the coverslips by dye exclusion. The video is briefly paused on the frame immediately after the application of strain to show the initial shape of the buckled tissue. Scale bar, 20 μm. Time is in mm:ss.

### Video 2

Buckling and flattening of a HaCaT epithelial tissue in response to a −35% compressive strain at high strain rate (500 %.s^-1^). Cell membranes are marked with CellMask (green), the medium is marked with Dextran Alexa647 (red) to allow for visualisation of the coverslips by dye exclusion. The video is briefly paused on the frame immediately after the application of strain to show the initial shape of the buckled tissue. Scale bar, 20 μm. Time is in mm:ss.

### Video 3

Buckling and partial flattening of an MDCK epithelial monolayer in response to a compressive strain of 50% applied at high strain rate (500 %.s^-1^). Cell membranes are marked with CellMask (green), the medium is marked with Dextran Alexa647 (red) to allow for visualisation of the coverslips by dye exclusion. The video is briefly paused on the frame immediately after the application of strain to show the initial shape of the buckled tissue. Scale bar, 20 μm. Time is in mm:ss.

### Video 4

Length adaptation and buckling of an MDCK epithelial monolayer in response to cycles of compressive strain applied at low strain rate (maximum strain: 80%). Cell membranes are marked with CellMask (green), the medium is marked with Dextran Alexa647 (red) to allow for visualisation of the coverslips by dye exclusion. Scale bar, 20 μm. Time is in mm:ss.

### Video 5

Buckling and partial flattening of an MDCK epithelial monolayer treated with 20 μM blebbistatin in response to application of 40% compressive strain applied at high strain rate. Cell membranes are marked with CellMask (green), the medium is marked with Dextran Alexa647 (red) to allow for visualisation of the coverslips by dye exclusion. Scale bar, 20 μm. Time is in mm:ss.

### Video 6

Dependence of flattening time on past strain history. Monolayers were subjected to the sequence of deformation shown in Fig. 4c. After an initial 6 minutes period of compressive strain, the tissue is returned to its initial length for Δt_1_ = 3 seconds and shortened again. Left: Tissue flattening in response to the initial cycle of compressive strain. Right: Faster tissue flattening in response to the second cycle of compressive strain. The video is paused briefly at 2 seconds and 15 seconds to allow comparison of the extent of flattening (see white dashed lines). Cell membranes are marked with CellMask (green), the medium is marked with Dextran Alexa647 (red) to allow for visualisation of the coverslips by dye exclusion. Scale bar, 20 μm. Time is in mm:ss.

### Video 7

Reversibility of change of flattening time due to strain history. MDCK monolayers were subjected to the sequence of deformation shown in Fig. 4c. Before a third cycle of compressive strain, the tissue is returned to its initial length for Δt_2_ = 6 minutes before the application of the third compression. Left: Tissue flattening in response to the initial cycle of compressive strain. Right: Tissue flattening in response to the third cycle of compressive strain. The flattening time is no longer distinguishable from the flattening time during the first cycle. The video is paused briefly at 2 seconds and 15 seconds to allow comparison of the extent of flattening (see white dashed lines). Cell membranes are marked with CellMask (green), the medium is marked with Dextran Alexa647 (red) to allow for visualisation of the coverslips by dye exclusion. Scale bar, 20 μm. Time is in mm:ss.

---

**Figure S1.**
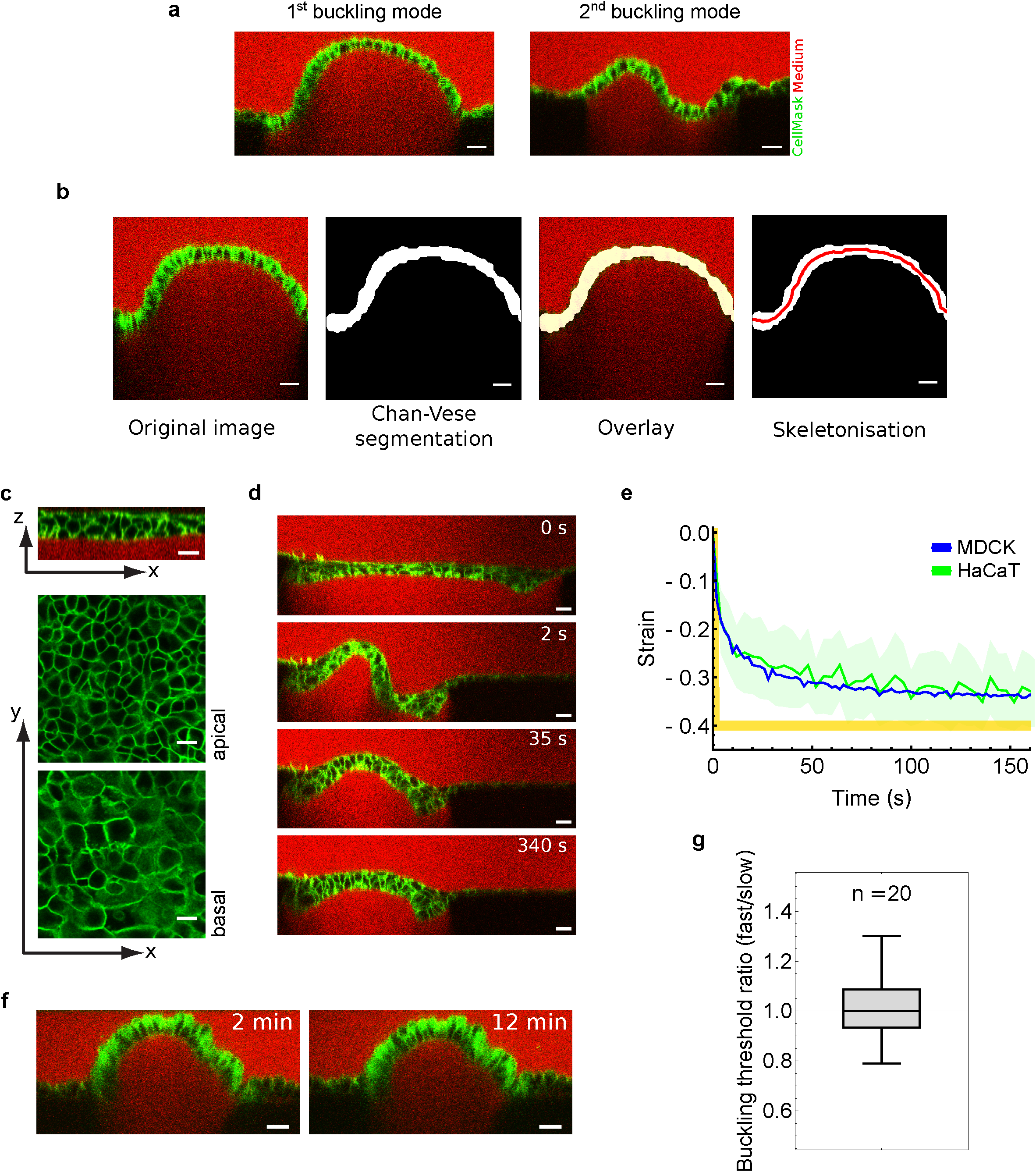
**a** – Profile view of the two tissue cross-sectional shape configurations observed immediately after application of compressive strain at high strain rate. Left: most frequently, a single arched shape was produced (68% of cases). Right: more complex shapes, reminiscent of the second mode of buckling, were produced in 32% of the cases. **b** – Automated video analysis pipeline for extraction of tissue contour length. From left to right, the cell membranes of the epithelial tissue are marked with CellMask (green) and imaged using scanning-laser confocal microscopy. The profile image is segmented using the Chan-Vese algorithm. The segmentation is then skeletonised to extract the contour length and tissue strain (see methods). **c** – Organisation of a suspended HaCaT tissue. Top: Profile view of the tissue showing multi-layering. Middle: single x-y confocal section of the apical surface of the tissue. Bottom: single x-y confocal section of the basal surface of the tissue. **d** – Time series of profile views of a suspended HaCaT epithelial tissue following application of compressive strain at high strain rate (500 %.s^-1^). Time is indicated in the top right corner. Scale bars, 20 μm. **e** – Temporal evolution of tissue strain in HaCaT epithelial tissues (green, mean ± SD) and MDCK epithelial monolayers (blue, mean ± SD) following application of compressive strain at high strain rate. The applied strain is shown in yellow. **f** – Stable fold induced by application of a strain larger than the buckling threshold in MDCK monolayers. Folds are stable for more than 10 minutes. Time is indicated in the top right corner. Scale bars, 20 μm. **g** – Ratio of the buckling threshold ε_b_ measured following a step of fast compressive strain over the buckling threshold ε_b_ measured following a slow ramp of compressive strain applied on the same sample, in a randomised order. The number of monolayers is indicated above the boxplot.

**Figure S2.**
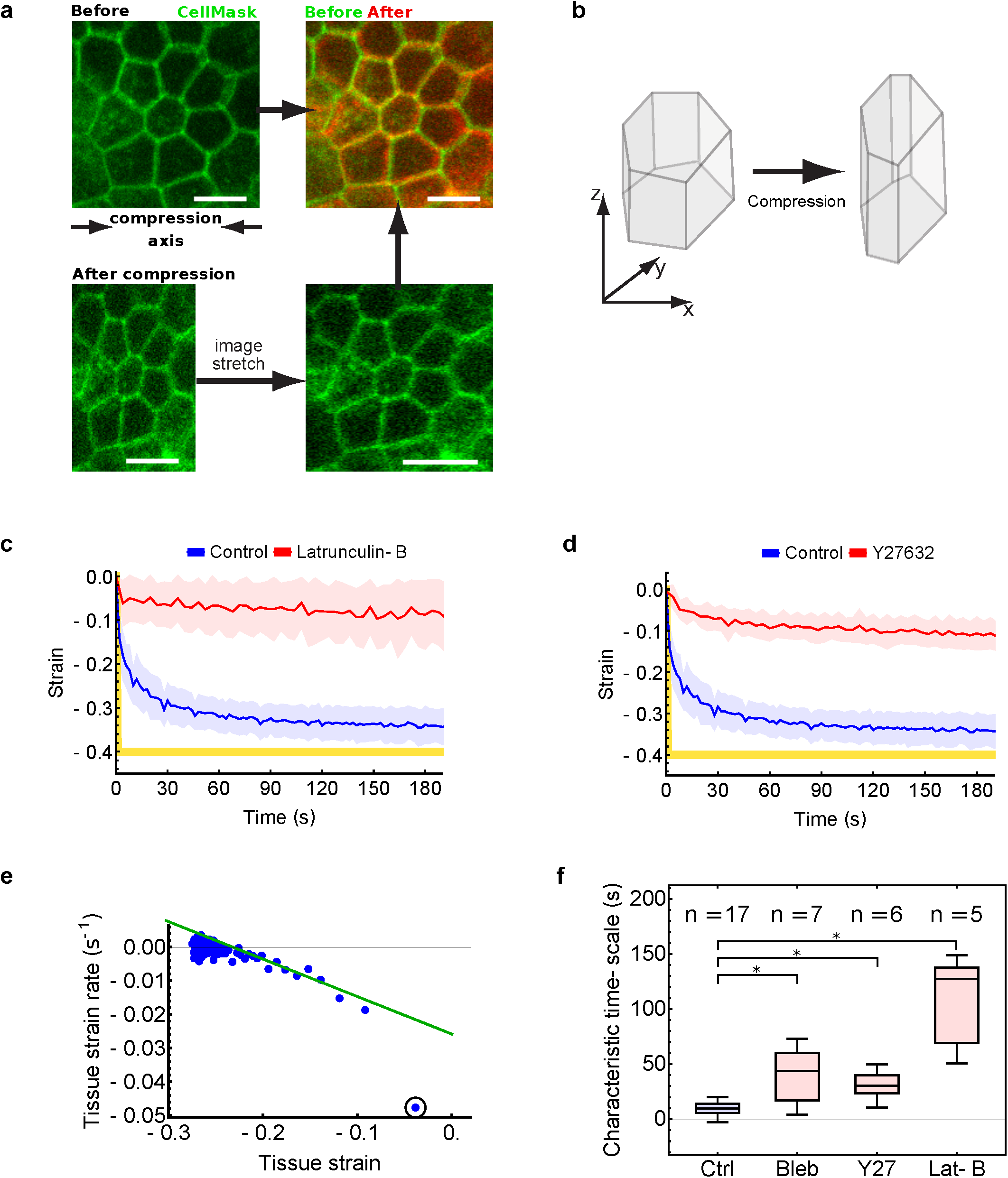
**a** – Tissue deformation after application of compressive strain is accounted for by isometric cellular deformations. Apical cell-cell junctions (CellMask membrane dye) from the same region of a MDCK monolayer are imaged before and after 30% compressive strain (left column). The image of the compressed tissue is then stretched numerically by 30% along the axis of experimentally applied deformation (bottom right). The overlay between this image and the image of the tissue before compression (top right) demonstrates that no topological changes occurred in the junctional network during compression. Scale bars, 10 μm. **b** – Diagram depicting the average cell shape change after application of 30% compressive strain. Cells reduce their length and increase their height to maintain a constant volume. **(c – d)** – Temporal evolution of tissue strain (mean ± SD) following compressive strain applied at high strain rate for control (blue) and treated tissues (red). Tissues were treated with Latrunculin B (**c**) or Y27632 (**d**). The device strain is shown in yellow. **e** – Extraction of a characteristic time-scale of the flattening process. Tissue strain rate was plotted against tissue strain following fast application of compression (blue dots). Strain rate is computed through a linear fit of 3 consecutive time-points from the evolution of strain over time (see example in Fig. 1d). Note that the first points (black circle) do not follow the linear regime and so were excluded from the fitting (green line); this is likely to be due to the power-law behaviour of MDCK monolayers described at short time-scales [38]. The characteristic time-scale of the flattening was defined as the negative reciprocal of the linear fit to this data. **f** – Characteristic time-scale of tissue flattening for tissues treated with a range of actomyosin contractility inhibitors. * denotes statistically significant difference, (p < 0.05). The number of monolayers examined is indicated above each box.

**Figure S3.**
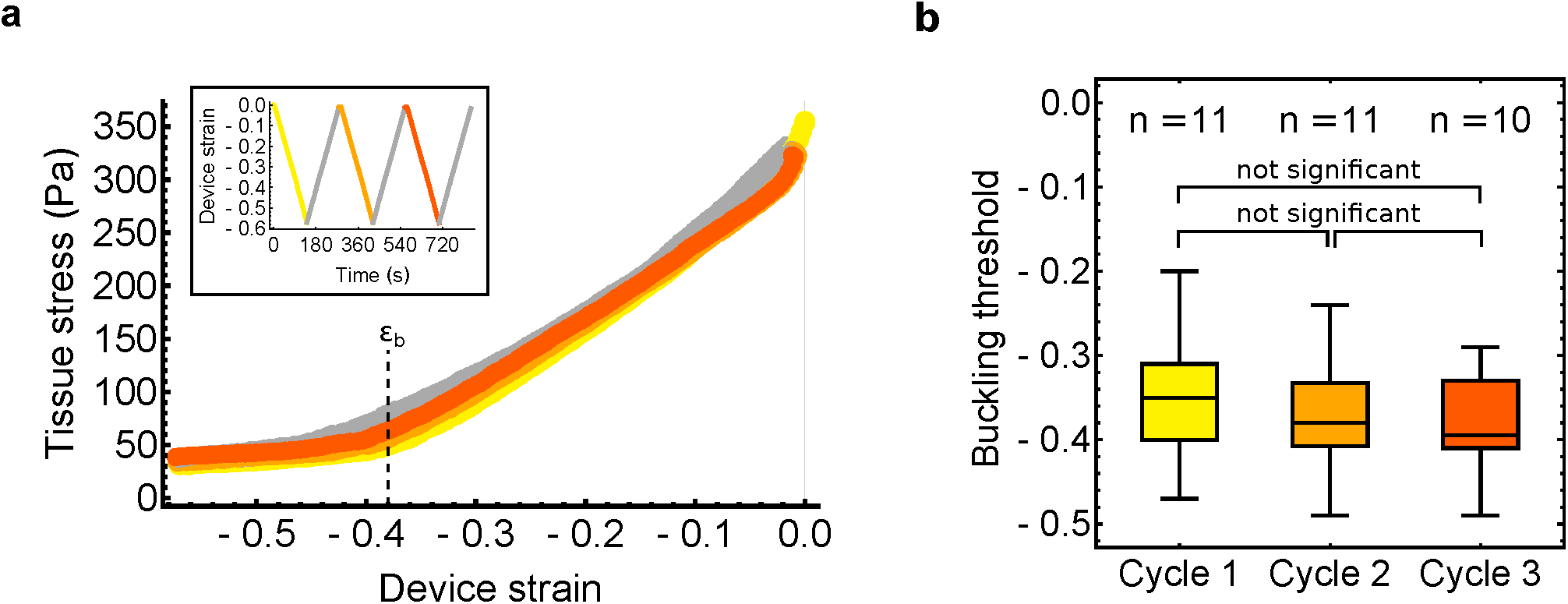
**a** – Tissue stress as a function of the applied device strain during deformation at low strain rate (0.5 %.s^-1^). Inset: Time course of the strain applied with the device. Three cycles of compressive strain were applied. Unloading is indicated in grey. The stress follows the same trend over 3 cycles of compressive strain shown in yellow (1^st^ cycle), orange (2^nd^ cycle) and dark orange (3^rd^ cycle). The dashed line marks the buckling threshold ε_b_. **b** – Comparison of the buckling threshold measured from the transition identified in stress-strain curves during each of the three cycles of applied compressive strain. There was no significant difference between the three cycles (p > 0.2). The number of monolayers examined in each condition is indicated above each box.

**Figure S4.**
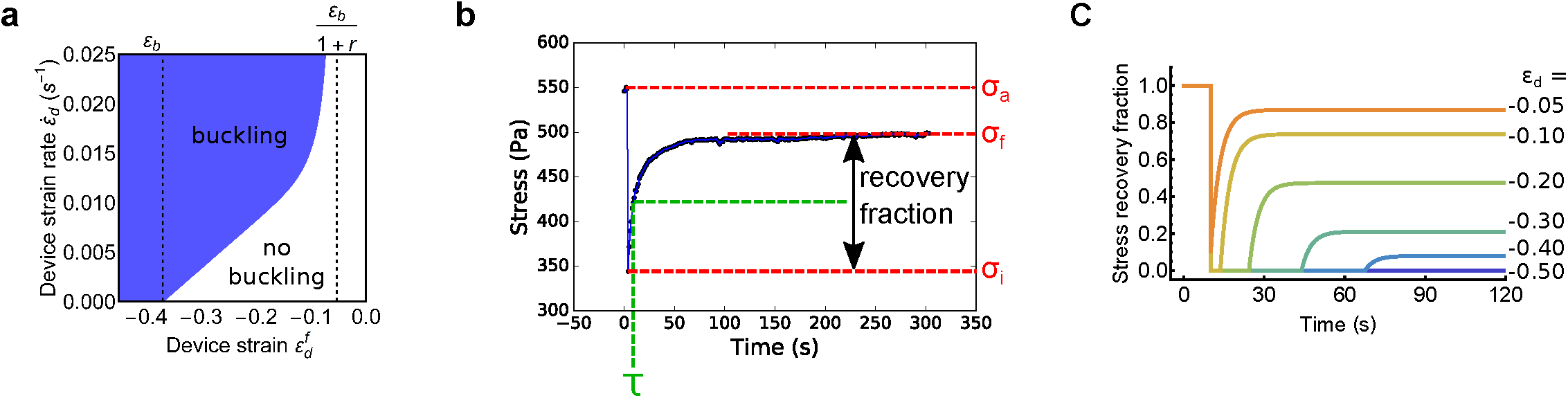
**a** – Phase diagram of buckling as a function of the device strain rate (y-axis) and the device strain (x-axis) predicted by the rheological model in Fig. 4a. (see Appendix 1 for details). The dashed lines delineate the buckling threshold ε_b_ (left) and the predicted smallest device strain which can induce a transient buckle (right). r is the ratio between the stiffnesses of the two springs in the model, Y/E. **b** – Representative temporal evolution of stress in response to low amplitude compressive strain applied at high strain rate. Tissue stress values (red) and the recovery time-scale (green) were extracted from the plot as shown and used to calculate the model parameters, as described in Appendix 2. **c** – Simulated tissue response to compressive strain of different magnitudes applied at high strain rate. The magnitude of device strain ε_d_ is shown on the right. Note the presence of a lag phase at zero stress corresponding to the duration over which the tissue is buckled. This lag phase is more pronounced than in experiments.

**Figure S5.**
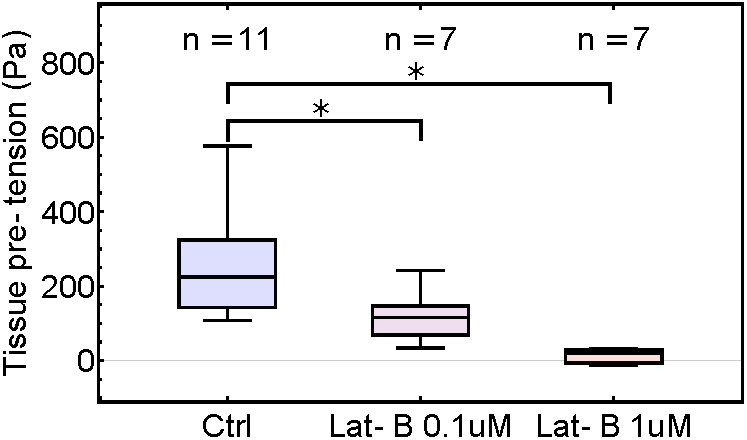
**a** – Tissue pre-tension measured for control and Latrunculin B treated samples (0.1 μM and 1 μM). * denotes statistically significant difference, p < 0.05. The number of monolayers examined in each condition is indicated above each box.

